# Glutamine metabolism modulates chondrocyte inflammatory response

**DOI:** 10.1101/2022.06.09.495504

**Authors:** Manoj Arra, Gaurav Swarnkar, Naga Suresh Adapala, Syeda Kanwal Naqvi, Lei Cai, Farooq Rai, Gabriel Mbalaviele, Robert Brophy, Yousef Abu-Amer

## Abstract

Osteoarthritis is the most common joint disease in the world with significant societal consequences, but lacks effective disease modifying interventions. The pathophysiology consists of a prominent inflammatory component that can be targeted to prevent cartilage degradation and structural defects. Intracellular metabolism has emerged as a culprit of the inflammatory response in chondrocytes, with both processes co-regulating each other. The role of glutamine metabolism in chondrocytes, especially in the context of inflammation, lacks a thorough understanding and is the focus of this work. We display that chondrocytes utilize glutamine for energy production and anabolic processes. Furthermore, we show that glutamine deprivation itself causes metabolic reprogramming and decreases the inflammatory response of chondrocytes through inhibition of NF-κB activity. Finally, we display that glutamine deprivation promotes autophagy and that ammonia is an inhibitor of autophagy. Overall, we identify a relationship between glutamine metabolism and inflammatory signaling and display the need for increased study of chondrocyte metabolic systems.

## Introduction

Joint disease afflicts millions of individuals around the world, though these conditions are greatly understudied. While many different diseases can affect the joints, osteoarthritis is the most common, affecting over 200 million individuals(Allen, Thoma, & Golightly, 2022). Significant advancements have been made in the treatment of classical inflammatory joint diseases, such as rheumatoid arthritis or psoriatic arthritis, due to identification of therapeutic targets and biomarkers(Palfreeman, McNamee, & McCann, 2013; Shams et al., 2021). Disease modifying compounds, especially with the advent of biologics, have revolutionized the management of these patients and allowed improved joint outcomes. Osteoarthritis (OA) remains a black sheep in this family of joint problems, lacking disease modifying interventions. OA can affect many different joints and presents in a variety of manners, likely contributing to the lack of clear disease pathophysiology or therapies, though inflammatory and biomechanical factors play a role, amongst other factors (Deveza & Loeser, 2018; Mobasheri & Batt, 2016). OA is characterized by joint degradation, with articular cartilage damage and joint space narrowing. Clinically, patients have increased pain and loss of joint mobility that can progress to significantly impair functionality. Interventions include use of NSAID’s and intra-articular steroid injections for pain relief, as well as joint replacement surgeries. However, there are currently no medications for preventing or reversing joint damage caused by osteoarthritis, and several clinical trials have failed(Hermann, Lambova, & Muller-Ladner, 2018). Furthermore, the disease is indolent and slowly progressing, often presenting with its classical symptoms long after joint degradation has begun and usually long after inciting factors such as joint injury have taken place (Blasioli & Kaplan, 2014; Roos & Arden, 2016). Clearly, there is a need for novel biomarkers, therapeutic targets, and overall understanding of disease pathophysiology.

Various groups have shown that inflammation is a driver of OA, even though it may not present like traditional inflammatory diseases such as rheumatoid arthritis (Arra, Swarnkar, Alippe, Mbalaviele, & Abu-Amer, 2022; Arra et al., 2020; Goldring & Otero, 2011). Joint inflammation causes cartilage-resident chondrocytes, as well as other joint infiltrating cells, to generate catabolic enzymes to promote an overall joint degradative state(Blasioli & Kaplan, 2014). Inflammatory stimuli activate signaling pathways such as the NF-κB pathway, which are important drivers of OA disease but have not been successfully targeted in OA(Arra et al., 2022; Arra et al., 2020; Catheline et al., 2021; Choi, Jo, Park, Kang, & Park, 2019). These stimuli can range from cytokines, to mechanical stress to inorganic particulate matter, making it difficult to target specific inflammatory mediators(Liu-Bryan & Terkeltaub, 2015; van den Bosch, 2019). Due to this, it is necessary to identify targetable cellular processes that modulate downstream inflammatory and catabolic activity in articular chondrocytes in response to a variety of stimuli.

Metabolic reprogramming is one such process that has gained interest in various cell types and disease state as a disease driver, and may also be important in OA (Chiellini, 2020; L. Zheng, Zhang, Sheng, & Mobasheri, 2021). Intracellular metabolism does far more than energy production and can modulate the inflammatory response of cells through regulation at various levels, ranging from epigenetics modifications to redox modulation(Gaber, Strehl, & Buttgereit, 2017; Lu & Wang, 2018). As such, chondrocyte intracellular metabolism has come into focus recently as a potential therapeutic target for modulating catabolic activity through regulation of inflammatory signaling pathways. Supporting this finding, we have shown recently that inflammatory stimuli alter the metabolism of chondrocyte, which can then regulate inflammatory responses (Arra et al., 2020).

Glucose metabolism has been fairly extensively studied in chondrocytes, though the role of other substrate pathways, such as fatty acid or amino acid metabolism, has been less well-studied. However, understanding the role of these other substrate pathways is critical since many metabolic pathways are interconnected and likely play a role in OA pathogenesis. In support of this claim, several groups have displayed recently that modulation of metabolic pathways can protect against OA and RA in animal models (Abboud et al., 2018; Coleman et al., 2018; Liu-Bryan, 2015; Ohashi et al., 2021; Shen et al., 2019). In humans, studies have shown that OA and RA joints have altered metabolite levels in the synovial fluid, though it is unclear if these changes are due to disease or drivers of disease(Akhbari et al., 2020; Kim et al., 2014; Zhai, 2019; K. Zheng et al., 2017). Finally, it has been displayed that systemic metabolic diseases such as obesity, diabetes and hypercholesterolemia (Baudart, Louati, Marcelli, Berenbaum, & Sellam, 2017) likely influence OA development, potentially through nutrient availability(Sellam & Berenbaum, 2013; Zhuo, Yang, Chen, & Wang, 2012). Based on these findings, it is probable that altered cell metabolism can be used not only as a biomarker of joint health but also as a therapeutic target. To help address some of the knowledge gaps in the field, we focused in this study on the role of glutamine in chondrocyte physiology and in response to inflammatory stimulation.

Glutamine is highly abundant throughout the body and is essential for many anabolic processes, but can also be utilized for energy production(Cruzat, Macedo Rogero, Noel Keane, Curi, & Newsholme, 2018). Some recent studies have elaborated the function of glutamine metabolism in chondrocyte physiology, though more understanding is required. One group demonstrated that inflammatory stimulation altered glutamine uptake and glutamate release in chondrocytes, with glutamate receptor involved in modulation of chondrocytes inflammatory response(Piepoli et al., 2009). Another group showed that glutamine metabolism is critical for regulating anabolic activity, glutathione production and epigenetic modifications in chondrocytes (Stegen et al., 2020). Furthermore, several studies have highlighted that glutamine levels are altered in synovial fluid of OA(Akhbari et al., 2020; Anderson et al., 2018). These studies highlight that glutamine is likely to be an important substrate for chondrocytes, both in healthy and disease states. In this study, we aim to characterize the role of glutamine metabolism in the inflammatory response of chondrocytes. We focus on NF-κB signaling as well as autophagy, both of which have been shown to be cooperatively important players in OA disease.

## Results

### Chondrocytes utilize glutamine for intracellular energy metabolism

We have previously displayed that chondrocytes under inflammatory conditions undergo metabolic reprogramming, with increased reliance upon glycolysis and decreased oxidative phosphorylation (Arra et al., 2020). Several studies have demonstrated that there is mitochondrial dysfunction with inflammatory stimulation(Arra et al., 2020; Lopez-Armada et al., 2006). During these conditions, there is decreased reliance upon glucose as a source of TCA cycle substrates, though other energy substrates such as glutamine can still fuel TCA activity to drive anabolic reactions (Martinez-Reyes & Chandel, 2020; Meiser et al., 2016). Thus, we seek to determine if chondrocytes can utilize glutamine to fuel anaplerotic TCA cycle activity. We observe that IL-1β stimulation alters the expression of glutamine and glutamate transports, as well as several key glutamine metabolic enzymes (Fig 1A-E).

**Fig 1.**
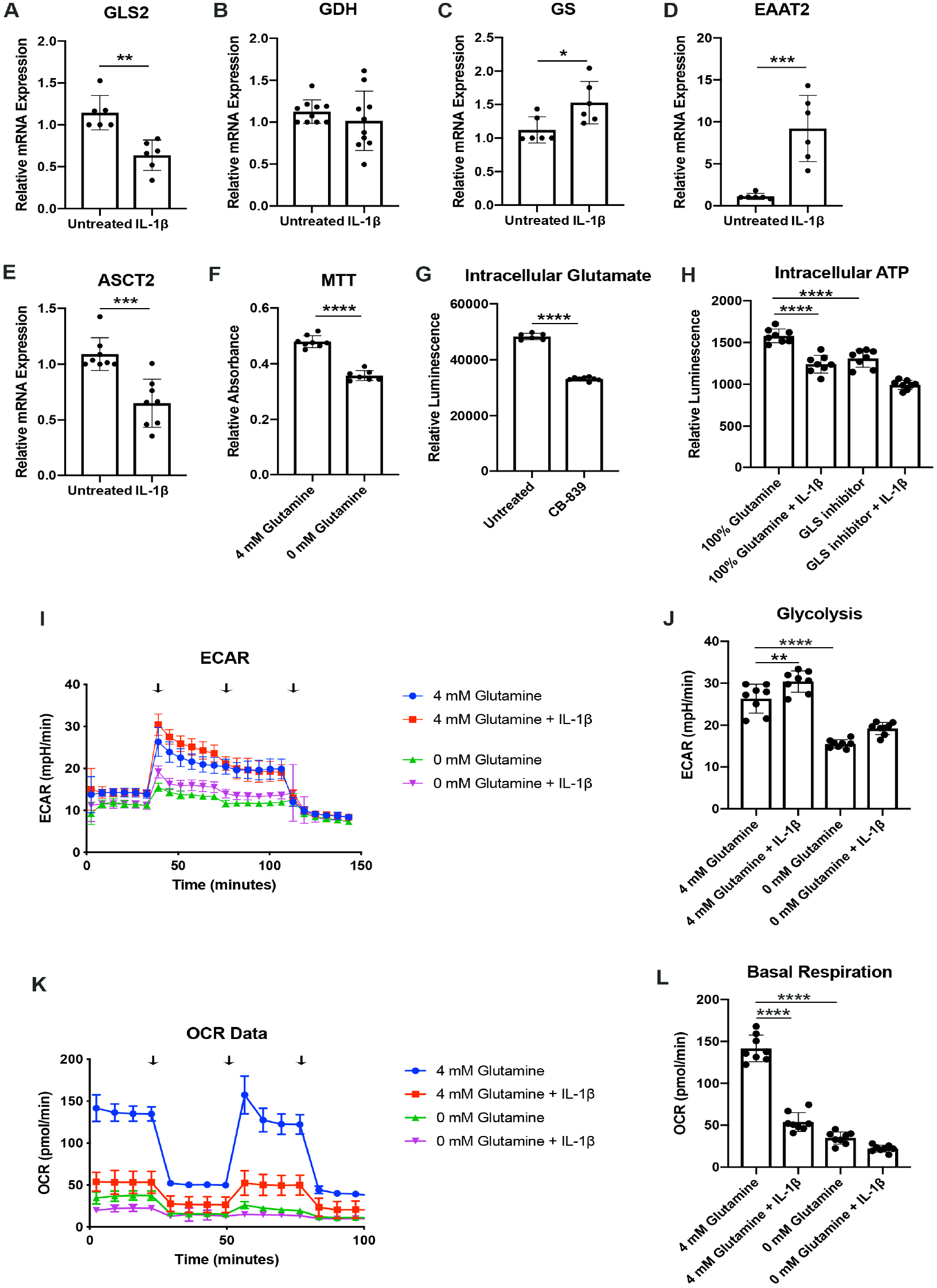
Chondrocytes rely upon glutamine for energy production. **(A-E)** Primary murine chondrocytes were treated with IL-1β (10 ng/mL) for 24 hours. Gene expression of *Gls2*, *Gdh*, *Gs*, *Eaat2* and *Asct2* were measured by qPCR. Results from n=6 independent biological samples. Unpaired student’s T test was performed. (A: **P=0.0011, B: P=0.375, C: *P=0.0227, D: ***P=0.0005, E: ***P=0.0003) **(F)** Primary murine chondrocytes were cultured in media with 4 mM glutamine and 0 mM glutamine under constant glucose conditions. After 24 hours, viability was measured by MTT assay. Results from n=6 samples from one representative experiment. Unpaired student’s T test was performed, ****P<0.0001. **(G)** Primary murine chondrocytes were treated with CB-839 (1 uM). Intracellular glutamate was measured by luminescent assay. Unpaired student’s T test was performed, ****P<0.0001. **(H)** Primary murine chondrocytes were treated with CB-839 and/or IL-1β for 24 hours. Intracellular ATP was measured by luminescent assay. Results from one representative experiment. One-way ANOVA was performed followed by Tukey’s multiple comparisons test. ****P<0.0001 **(I-L)** Primary sternal chondrocytes were cultured in media containing glutamine or media without glutamine for 24 hours. Cells were then treated with IL-1β (10 ng/mL) for 24 hours. All values were normalized to cell viability of treatments relative to untreated cells as measured by MTT assay. (**I-J**) ECAR measurement in glycolysis stress test (Injection 1: No treatment, Injection 2: Glucose, Injection 3: Oligomycin, Injection 4: 2-DG) or (**K-L**) OCR measurement in MitoStress test (Injection 1: No treatment, Injection 2: Oligomycin, Injection 3: FCCP, Injection 4: Antimycin A/Rotenone) was performed on Seahorse Instrument. Measurements were performed every 6 minutes with 8 replicates per timepoint for each condition. Arrows represent injections timepoints. Graphs shown in Fig 1J and 1L are from a single timepoint. One-way ANOVA was performed followed by Tukey’s multiple comparisons test. J:**P=0.0077, ****P<0.0001, L:****P<0.0001.

However, enzyme expression may not necessarily reflect glutamine utilization. To determine this, we measured viability of chondrocytes in the presence or absence of glutamine. Culturing chondrocytes in glutamine free media led to a slight decrease in viability, suggesting that chondrocytes require glutamine for energy production (Fig 1F). Since the first step of glutamine metabolism and entry to TCA cycles is conversion to glutamate, a process catalyzed by glutaminase (GLS), we utilized a GLS inhibitor, CB-839, which mimics the effect of glutamine deprivation on cell metabolism by preventing glutamate generation. We confirmed that GLS inhibition significantly reduced intracellular glutamate to levels approaching that of glutamine deprivation (Fig 1G). We also observed a decrease in ATP levels with GLS inhibition, further confirming that chondrocytes do utilize glutamine for energy production (Fig 1H). Given that chondrocytes clearly rely upon glutamine, we then performed Seahorse analysis to confirm that chondrocytes utilized glutamine for metabolism and energy production. We note that IL-1β stimulation increases glycolysis and causes a dramatic decrease in OxPhos, likely via mitochondrial dysfunction (Fig 1I-L). We observed that glutamine deprivation led to a decrease in both extracellular acidification rate (ECAR) and oxygen consumption rate (OCR), with a more dramatic effect on OCR, likely due to contribution of glutamine to TCA anaplerotic activity (Fig 1I-L). We observe similar effects with GLS inhibition, indicating that glutamine breakdown by GLS is an essential step for energy metabolism (Figure 1 Figure Supplement 1A-D).

We sought to determine if human OA cartilage also displays altered expression of glutamine metabolic enzymes. We noted that human OA cartilage display increased expression of various glutamine metabolic enzymes (Figure 1 Figure Supplement 1 E-H), potentially suggesting increased glutamine metabolism. We validate that OA cartilage is more catabolic and inflammatory through measurement of NF-κB Inhibitor Zeta (*Nfkbiz)* and Matrix metalloprotease 3 (*Mmp3)* expression, respectively (Fig I-J). We also note that IL-1β stimulation of human chondrocytes isolated from knee cartilage caused some glutamine metabolic enzyme changes, though less significantly than OA chondrocytes (Figure 1 Figure Supplement 1K-M), indicating that metabolic changes in human cells may be a chronic change or in response to other inflammatory stimuli.

### Glutamine deprivation causes metabolic reprogramming to inhibit glycolysis

We were surprised to observe that glutamine deprivation of chondrocytes was able to cause a reduction in both glycolysis and oxidative phosphorylation, given that glutamine primarily supplies TCA cycle activity(Yoo, Yu, Sung, & Han, 2020). We sought to determine what impact glutamine deprivation may have on systems such as glycolysis which tend to primarily utilize glucose, especially in the context of inflammation. Furthermore, we have previously displayed that metabolic reprogramming towards increased glycolysis induced by IL-1β can promote catabolic activity and OA disease(Arra et al., 2020).

We observe that glutamine deprivation itself was able to induce metabolic reprogramming that supports TCA activity. We noted increased expression of glutaminase *(Gls)* with glutamine deprivation (Figure 2 Supplement 2A) and slight increase in some TCA cycles enzyme expression such as malate dehydrogenase (*Mdh)* and Succinate dehydrogenase subunit A (*Sdha)*, with insignificant changes in others (Figure 2 Supplement 2B-E). However, glutamine deprivation inhibited the expression of various glycolytic and pentose phosphate pathway (PPP) enzymes (Fig 2A-C). Furthermore, we observe that glutamine deprivation can prevent many of the glycolytic and PPP enzymes metabolic changes observed with IL-1β stimulation (Fig 2A-C). This is further supported by the finding that glutamine deprivation can reduce lactate production by chondrocytes, a marker of glycolytic activity (Figure 2 Supplement 2F). Based on our findings, it appears that glutamine deprivation supports TCA cycle activity, but inhibits glycolysis and PPP. This may be a compensatory mechanism utilized by cells in the absence of glutamine to sustain ATP production via utilization of the energy favorable TCA cycle, as well as anabolic activity.

**Fig 2.**
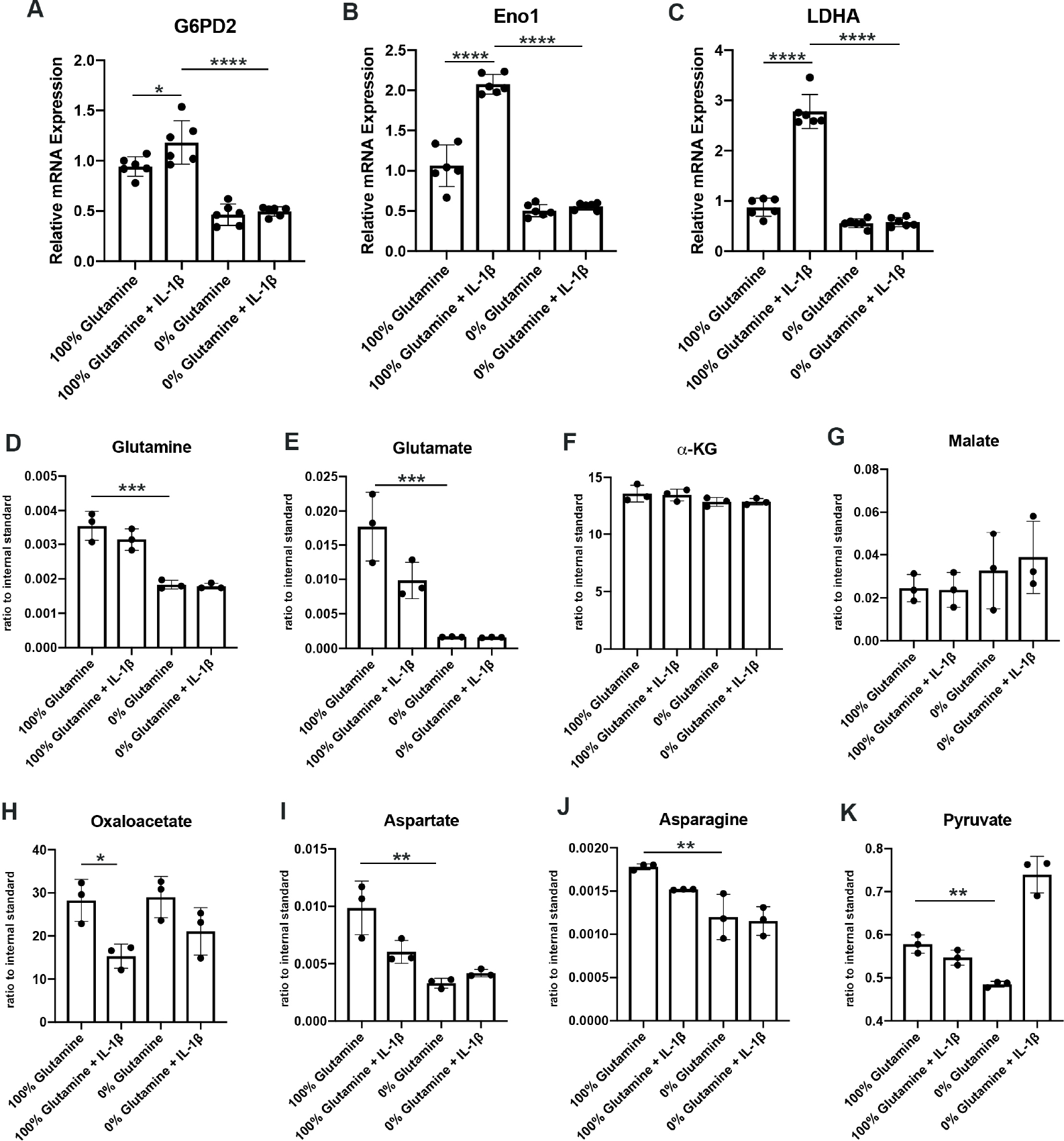
Glutamine deprivation causes metabolic reprogramming to inhibit glycolysis. **(A-C)** Primary murine chondrocytes were cultured in media containing 100% glutamine or 0% glutamine for 24 hours. Cells were then treated with IL-1β (10 ng/mL) for 24 hours. Gene expression of *G6pd2*, *Eno1* and *Ldha* were measured by qPCR from n=6 replicates. One-way ANOVA was performed followed by Tukey’s multiple comparisons test. A: *P=0.0242, ****P<0.0001, B: ****P<0.0001, C: ****P<0.0001 **(D-K)** Under similar conditions, metabolite levels were measured by LC-MS with n=3 replicates. One-way ANOVA was performed followed by Tukey’s multiple comparisons test. D: ***P=0.0003, E:***P=0.0006, F: P>0.05, G: P>0.05, H:*P=0.036, I: **P=0.0012, J:**P=0.0079, K: **P=0.009.

We then performed some targeted proteomics in the context of glutamine deprivation (Fig 2D-E) and noted that glutamine deprivation does not reduce levels of TCA metabolites such as α-KG, malate, and oxaloacetate, but does reduce levels of pyruvate generated from glycolysis (Fig 2D-K). However, we did note that glutamine is in fact required for the production of various downstream substrates, such as asparagine and aspartate (Fig 2I-J). Given that glutamine deprivation reduced OxPhos activity and ATP levels, but did not reduce TCA metabolite levels, chondrocytes may be able to generate TCA metabolites by utilization of other anaplerotic processes. These systems can generate metabolites but usually do not generate energy. As an example, we noted increased expression of *Psat1* with glutamine deprivation (Figure 2 Supplement 2G), which has recently been displayed to be one source of glucose-based α-KG to fuel TCA cycle (Hwang et al., 2016).

### Glutamine metabolism by glutaminase contributes to inflammatory gene expression

We have previously displayed that altered metabolism can regulate the inflammatory and catabolic response of chondrocytes. Since the metabolic changes induced by glutamine deprivation opposed the metabolic changes we observe with IL-1β stimulation, we suspected that glutamine may also play a role in modulating the inflammatory response induced by IL-1β.

To determine the role of glutamine metabolism in the inflammatory response, we cultured chondrocytes in media with and without glutamine under sufficient glucose conditions and treated them with IL-1β. We observed that glutamine deprivation led to a decrease in inflammatory and catabolic gene expression in response to IL-1β, with a reduction in expression of genes such as Interleukin 6 (*Il6)* and Matrix metalloprotease 13 (*Mmp13)* (Fig 3A-B). We then sought to determine the mechanism by which glutamine can regulate the inflammatory response. We measured NF-κB activity since it is the principle inflammatory response pathway to IL-1β that we have previously demonstrated is important for OA development(Arra et al., 2022; Arra et al., 2020). We observe using chondrocytes derived from p65-luciferase reporter mice that glutamine deprivation dose dependently inhibits NF-κB activation, as measured by luciferase activity (Fig 3C). It has also previously been displayed that IκB-ζ is another critical pro-inflammatory mediator of NF-κB activity in chondrocytes treated with IL-1β (Arra et al., 2022; Choi, MaruYama, Chun, & Park, 2018). We observe that glutamine deprivation leads to a decrease in IκB-ζ protein expression and stability, at least partially due to inhibition of NF-κB activity (Fig 3D). Our earlier work has also shown that IκB-ζ is a redox sensitive protein that is stabilized by oxidative stressors from metabolic sources such as LDHA and the mitochondria, so we measured ROS levels in the absence of glutamine in response to IL-1β stimulation. We surprisingly observed decreased ROS production in the absence of glutamine (Fig 3E), likely due to decreased inflammatory response and a reduction in pro-oxidative metabolic changes we have previously characterized.

**Fig 3.**
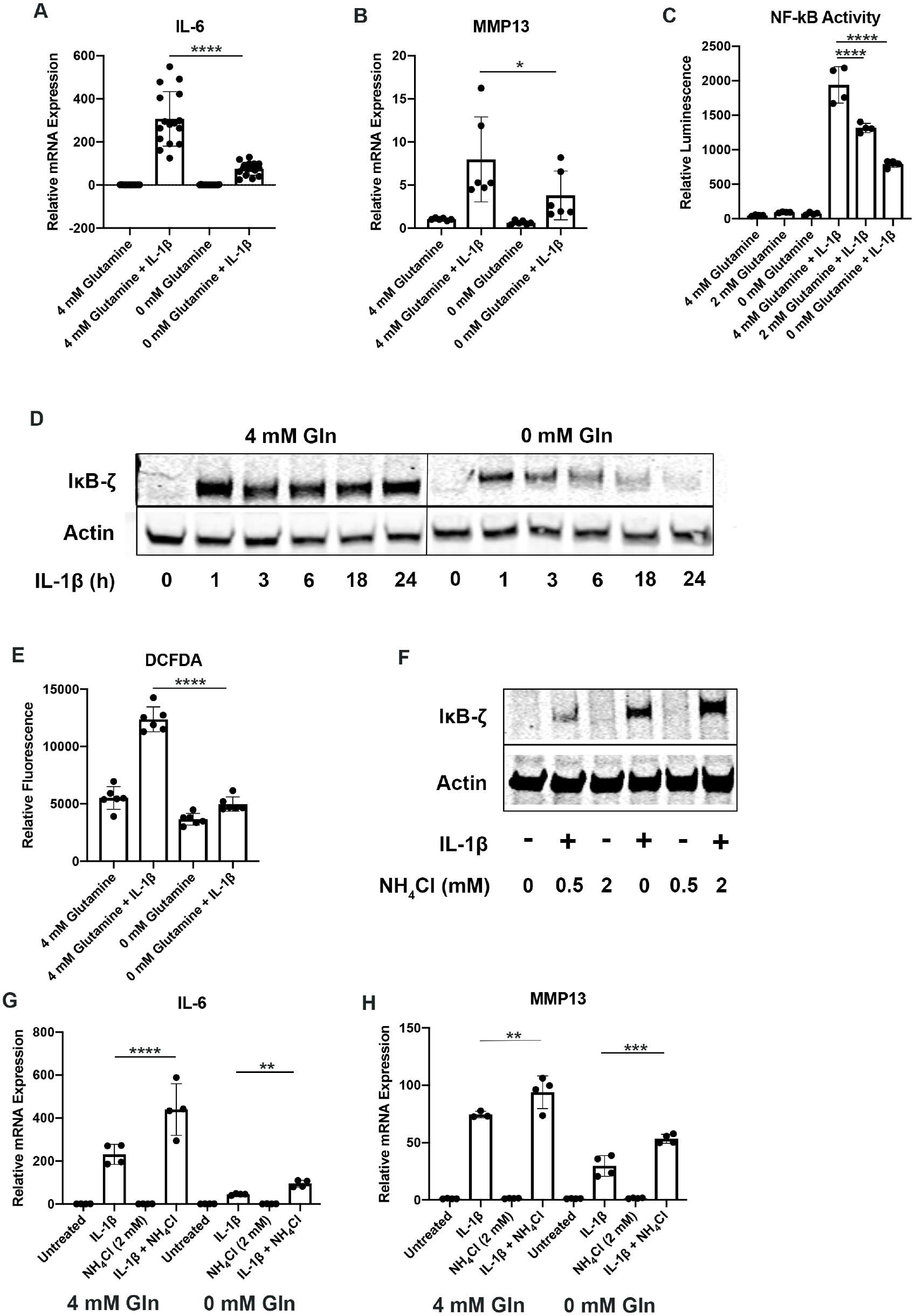
Glutamine deprivation inhibits the inflammatory response. **(A-B)** Primary murine chondrocytes were cultured in media containing 100% glutamine or 0% glutamine for 24 hours. Cells were then treated with IL-1β (10 ng/mL) for 24 hours. Gene expression of *Il6* and *Mmp13* were measured by qPCR. One-way ANOVA was performed followed by Tukey’s multiple comparisons test. A:****P<0.0001, B: *P=0.0489 **(C)** Primary murine chondrocytes were isolated from NF-κB-luciferase reporter mice. Chondrocytes were then cultured in media containing 4 mM, 2 mM or 0 mM glutamine for 24 hours. Cells were then treated with IL-1β for 24 hours. NF-κB activity was measured by luciferase assay. One-way ANOVA was performed followed by Tukey’s multiple comparisons test. ****P<0.0001. **(D)** Primary murine chondrocytes were cultured in media containing 100% glutamine or 0% glutamine for 24 hours. Cells were treated with IL-1β for the indicated timepoints. IκB-ζ protein was measured by immunoblotting, with actin used as housekeeping. Image displays representative experiment. **(E)** Primary murine chondrocytes were cultured in media containing 100% glutamine or 0% glutamine for 24 hours. Cells were then treated with IL-1β (10 ng/mL) for 24 hours. ROS levels were measured by DCFDA assay using microplate reader. One-way ANOVA was performed followed by Tukey’s multiple comparisons test. ****P<0.0001. **(F)** Primary chondrocytes were cultured in media containing glutamine and supplemented with ammonium chloride at the indicated concentrations for 24 hours in the presence of IL-1β. IκB-ζ protein was measured by immunoblotting. **(G-H)** Primary chondrocytes were cultured in media containing 4 mM or 0 mM glutamine for 6 hours. Cells were then supplemented with or without 2 mM ammonium chloride. IL-1β stimulation was performed for 24 hours. Gene expression of *Il6* and *Mmp13* was measured by qPCR. One-way ANOVA was performed followed by Tukey’s multiple comparisons test. G: ****P<0.0001, **P=0.0065, H: **P=, ***P= 0.0096, 0.0005.

We then interrogated if glutaminase inhibition can modulate the inflammatory response similar to glutamine deprivation. We observed that glutaminase inhibition by CB-839 was also effective at decreasing the inflammatory response, suggesting that glutamine to glutamate conversion is important for the inflammatory response (Figure 3 Supplement 3A). We also observed that glutaminase inhibition was also able to potently decrease IκB-ζ protein expression (Figure 3 Supplement 3B), indicating that glutaminolysis contributes to IκB-ζ-mediated gene expression. Furthermore, GLS inhibition reduced NF-κB activation (Figure 3 Supplement 3C).

The glutaminolysis reaction performed by GLS generates glutamate and free ammonia from glutamine(Yoo et al., 2020). Glutamate can then be converted to a-ketoglutarate, glutathione or undergo transaminase reactions. On the other hand, ammonia is a reactive species, often viewed as a waste product, and can be incorporated into amino acids or urea for its removal(Kurmi & Haigis, 2020; Spinelli et al., 2017). We sought to determine if glutamate or ammonia generated by GLS can regulate the inflammatory response of chondrocytes. We supplemented chondrocytes with ammonia or glutamate and measured the inflammatory response. We observed that ammonia supplementation was pro-inflammatory, activating NF-κB and increasing IκB-ζ gene expression in the setting of IL-1β stimulation (Fig 3F, Figure 3 Supplement 3D). It also increased expression of inflammatory and catabolic genes (Fig 3G-H). Ammonia supplementation also partially rescued inflammatory gene expression under glutamine deprivation conditions and IκB-ζ protein expression. Glutamate supplementation did not have a significant impact on NF-κB activation (Figure 3 Supplement 3D), and was unable to rescue IκB-ζ expression (Figure 3 Supplement 3E). This finding suggests that ammonia generation from glutamine metabolism may be involved in promoting inflammation.

Given that we did not observe an increase in inflammation with glutamate supplementation, we then tested the efficacy of GDH inhibitor, EGCG, which blocks the conversion of glutamate to α-KG(Li et al., 2006), a process that also releases an ammonia group and is critical for the generation of pro-inflammatory downstream metabolites such as succinate. We note that EGCG treatment also slightly reduced the inflammatory response represented by *Il6* expression, though less potently than glutamine deprivation (Figure 3 Supplement 3F).

### Glutamine deprivation activates autophagy

Since it is well known that nutrient deprivation can induce autophagy(Russell, Yuan, & Guan, 2014), we sought to determine what impact glutamine deprivation would have on chondrocyte autophagy processes, especially in the context of inflammation. We note that IL-1β stimulation of chondrocytes leads to a decrease in autophagy, as noted by accumulation of p62 protein. We then observe that there is an upregulation of autophagy with glutamine deprivation, as indicated by a decrease in p62 protein, which is often an indication of autophagy progression as p62 is degraded by autophagosomes (Fig 4A). We also note a significant decrease in LC3 protein levels with glutamine deprivation due to increased consumption of LC3 through autophagy. We validated this through chloroquine treatment, an inhibitor of autophagy, which can rescue LC3 and p62 levels in glutamine deprivation conditions, indicating that LC3 and p62 are being processed by autophagy processes. We validated these findings by immunofluorescence, which displayed that chloroquine treatment led to far greater increase in LC3-positive punctate in cells under glutamine deprivation conditions compared to glutamine replete conditions (Fig 4B). We confirm that findings are due to protein processing since we do not observe similar changes at the gene expression level (Figure 4 Supplement 4A-B). Interestingly, we note that glutamine deprivation also leads to a transient decrease in gene expression of Microtubule-associated protein 1A light chain 3b (*Lc3b)* and sequestosome (*p62)* at less than 24 hours, which recovers at the 24 hour timepoint (Figure 4 Supplement 4C-D). We also observe that LC3 protein levels start to decrease rapidly with glutamine deprivation but p62 levels do not decrease until 48 hours (Figure 4 Supplement 4E). We observed similar effects with GLS inhibition by CB-839 (Figure 4 Supplement 4F).

**Fig 4.**
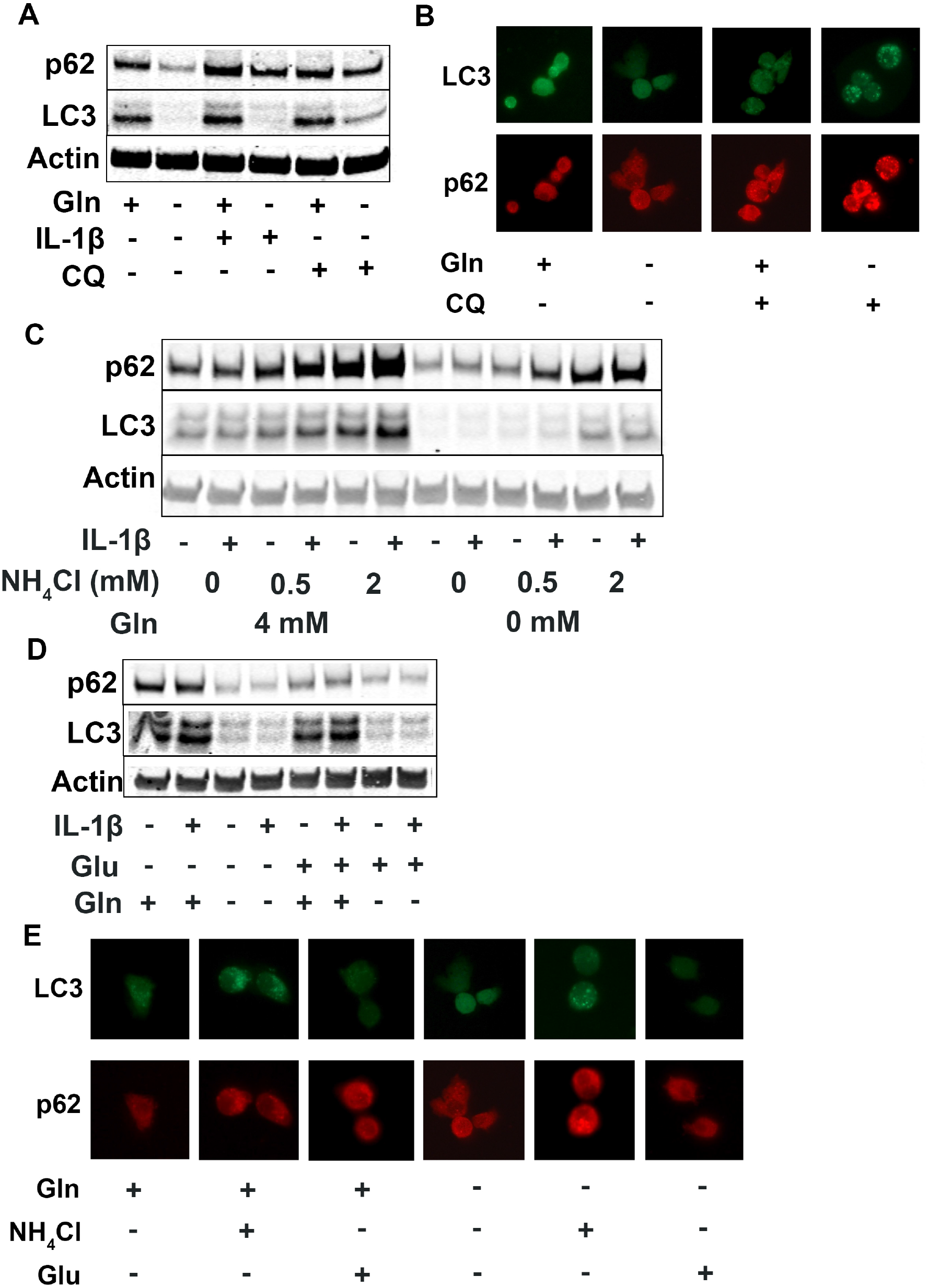
Glutamine deprivation promotes autophagy and ammonia inhibits autophagy. **(A)** Primary murine chondrocytes were cultured in media containing 100% glutamine or 0% glutamine for 24 hours. Cells were then treated with IL-1β (10 ng/mL) in the presence or absence of chloroquine (10 μM) for 24 hours. Protein expression of p62 and LC3-II were measured by immunoblotting. **(B)** Primary murine chondrocytes were plated on coated cover slips cultured in glutamine containing or glutamine free media for 12 hours. Cells were treated with chloroquine (10 μM) for 6 hours. Cells were fixed with 4% formaldehyde in PBS and immunofluorescence (IF) was performed for LC3B and p62. Cells were mounted on slides and imaged with representative images displayed. **(C)** Primary chondrocytes were cultured in media containing 4 mM or 0 mM glutamine. Cells were supplemented with ammonium chloride at the indicated concentrations. After 6 hours, cells were treated with IL-1β (10 ng/mL) for 24 hours. Immunoblotting was performed for p62 and LC3B to display autophagosome processing. Image displays representative experiment. (D) Primary chondrocytes were cultured in media containing 4 mM or 0 mM glutamine. Cells were supplemented with glutamate (200 μM). After 6 hours, cells were treated with IL-1β (10 ng/mL) for 24 hours. Immunoblotting was performed for p62 and LC3b. Image displays representative experiment. **(E)** Primary murine chondrocytes were plated on coated cover slips cultured in glutamine containing or glutamine free media for 12 hours. Cells were supplemented with ammonium chloride (2 mM) or glutamate (200 μM). Cells were fixed with 4% formaldehyde and IF was performed for LC3B and p62. Cells were mounted on slides and imaged with representative images to display autophagosome punctate.

We then supplemented chondrocytes deprived of glutamine with glutamate or ammonia and measured levels of LC3 and p62 to determine how glutaminolysis affects autophagy. We noted ammonia supplementation was able to inhibit autophagy and reverse the effect of glutamine deprivation on LC3 and p62 expression, similar to chloroquine treatment (Fig 4C, Figure 4 Supplement 4G). Ammonia treatment led to an increase in LC3b and p62, likely through blockade of autophageosome-lysosome fusion. We also observe that glutamate supplementation appeared to increase autophagy, as noted by a decrease in p62 levels (Fig 4D, Figure 4 Supplement 4G). These findings were confirmed through increased number of LC3 positive punctate in ammonia treated cells but not in glutamate treated cells, indicating opposing effects of ammonia and glutamate on autophagy (Fig 4E).

One of the major cell stress response factors involved in regulating metabolism and autophagy is Activating transcription factor 4 (ATF4), which can modulate intracellular metabolism(B’Chir et al., 2013; O’Leary et al., 2020; Stegen et al., 2022). Mechanistically, we observe that glutamine deprivation is able to increase ATF4 protein expression, which is reduced with IL-1β stimulation (Figure 4 Supplement 4H-I). We note that at around 12 hours of glutamine deprivation, *Atf4* expression increases (Figure 4 Supplement 4J). ATF4 activation is a well-known response system to amino acid deprivation, and it is known to be a driver of autophagy processes(Jin, Hong, Kim, Jang, & Park, 2021; Ye et al., 2010). ATF4 is also a mediator of metabolic reprogramming, which we observe with glutamine deprivation and IL-1β stimulation. Suspecting that ATF4 may be important in OA, we note that OA mouse cartilage (MLI) has decreased expression of ATF4, mimicking the effect of IL-1β stimulation (Figure 4 Supplement 4K). Based on these results, it is predicted that ATF4 may be an anti-inflammatory factor and will be the focus of future work.

### mTOR2 but not mTOR1 is activated by glutamine deprivation

Since mTOR signaling is another critical factor connecting metabolism and autophagy, we interrogated mTOR activation in the setting of glutamine deprivation and inflammation. mTOR activation has been shown to be a driver of metabolic changes, especially for glycolytic pathways(Linke, Fritsch, Sukhbaatar, Hengstschlager, & Weichhart, 2017; Magaway, Kim, & Jacinto, 2019). Glutamine deprivation decreased glycolysis and glycolytic enzyme expression, hence we suspected that mTOR modulation may be involved. We note that mTOR1 and mTOR2 activity is increased with IL-1β stimulation as measured by increased phosphorylation of S6 riboprotein and phosphorylation of AKT-473 (Fig 5A-B), supporting the inhibition in autophagy we previously measured. With glutamine deprivation, we noticed a decrease in mTOR1 activity with glutamine deprivation, as measured by phosphorylation of S6 ribosomal protein but an increase in mTOR2 activity, as measured by increased AKT-473 phosphorylation (Fig 5A-B).

**Fig 5.**
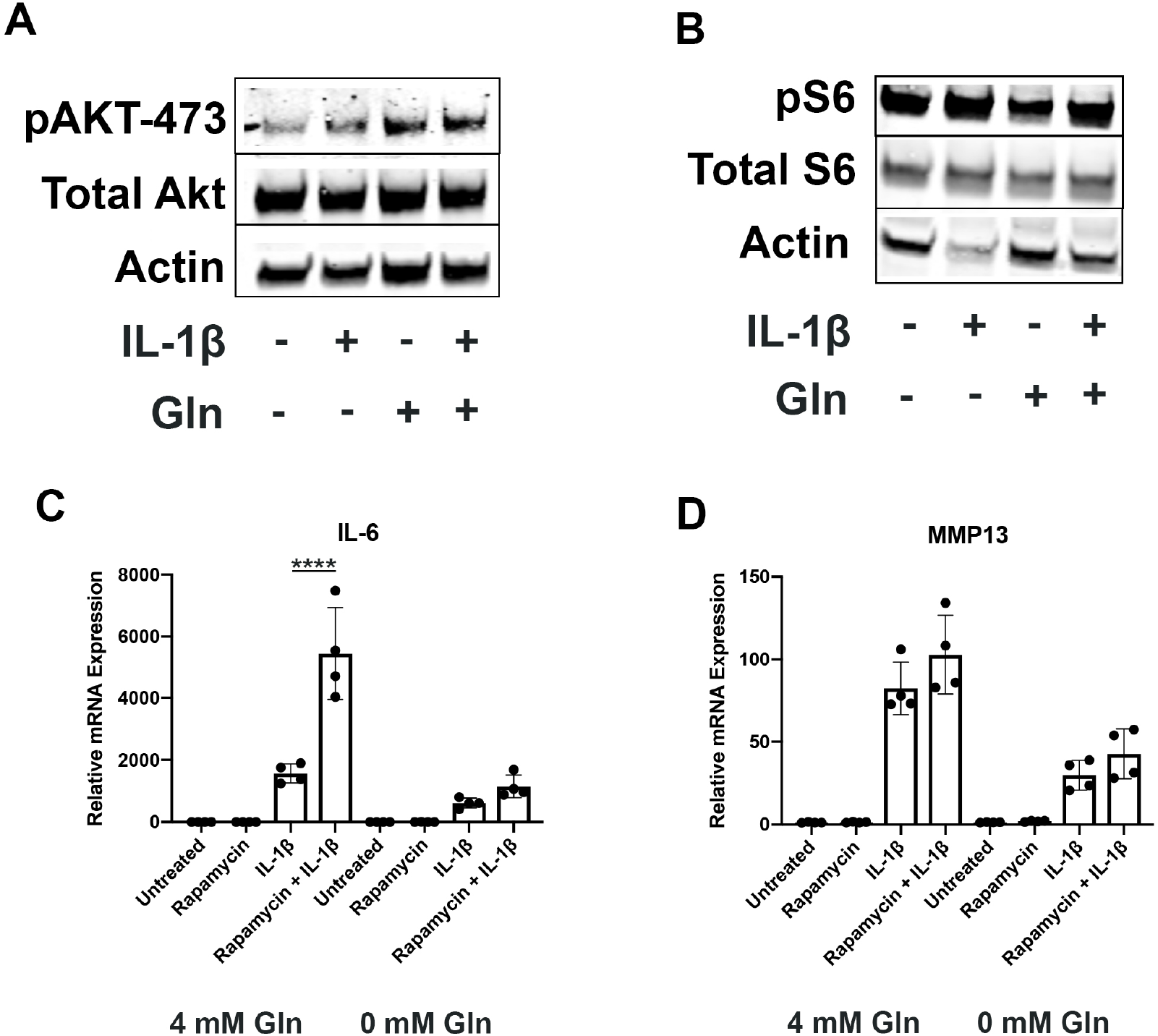
Glutamine deprivation modulates mTOR activation. **(A)** Primary murine chondrocytes were cultured in media containing 100% glutamine or 0% glutamine for 24 hours. Cells were then treated with IL-1β (10 ng/mL). After 24 hours, lysates were collected and immunoblotting was performed for pAKT and total Akt. **(B)** Under similar conditions, immunoblotting was performed for pS6 and total S6. **(C-D)** Primary murine chondrocytes were cultured in media containing 100% glutamine or 0% glutamine for 24 hours. Cells were then treated with IL-1β (10 ng/mL) in the presence or absence of rapamycin 50 nM for 24 hours. Gene expression of *Il6* and *Mmp13* was measured by qPCR. Results from one representative experiment. One-way ANOVA was performed followed by Tukey’s multiple comparisons test. C:****P<0.0001.

We then utilized rapamycin, an mTOR1 inhibitor, to interrogate the role of mTOR in the inflammatory response and glutamine metabolic gene expression changes. We validate that rapamycin can block mTOR1 activation through complete abrogation of phospho-S6 expression (Figure 5 Supplement 5A). We display that rapamycin treatment can reverse some metabolic changes induced by IL-1β, such as increased glycolytic enzyme expression using LDHA as a representative gene (Figure 5 Supplement 5B). We also note that rapamycin treatment is able to upregulate expression of Glutamine synthase (*Gs)* and *Gls*, but did not affect glutamic-oxaloacetic transaminase 2 (*Got2)* expression (Figure 5 Supplement 5C-E). However, under glutamine deprivation conditions, rapamycin treatment only affected *Gls* expression, but not *Gs* or *Got2*. We then measured the impact of mTOR1 inhibition by rapamycin on the inflammatory response, and surprisingly observed that rapamycin increases the expression of inflammatory and catabolic genes (Fig 5C-D). It also reverses some of the inflammatory inhibition induced by glutamine deprivation, suggesting an important role for mTOR1 in the anti-inflammatory effect of glutamine deprivation.

## Discussion

This work displays that glutamine utilization by chondrocytes is important for their physiology and inflammatory response, and may be an important player in OA. We show that chondrocytes utilize glutamine for energy production even when glucose is abundant, indicating that chondrocytes rely upon multiple substrates for metabolism. However, we observe that glutamine deprivation is able to decrease chondrocyte glycolysis and oxidative phosphorylation, which was surprising since prior groups have mainly described glutamine as fueling the TCA cycle in the mitochondria. This finding was explained when we noted that glutamine deprivation caused metabolic reprogramming to inhibit glycolytic activity by significantly decreasing expression of glycolytic enzymes, but maintained TCA metabolites. This was a surprising new interaction between glutamine and glucose metabolism which may be a compensatory mechanism that forces chondrocytes to rely upon other energy sources such as fatty acid oxidation to maintain anabolic processes. In addition, this reprogramming caused by glutamine deprivation may actually be protective in the context of inflammation, as will be discussed further. Future work utilizing an untargeted metabolomics approach will be useful for understanding the contribution and interactions of various substrate pathways to chondrocyte metabolism and physiology.

From an inflammatory standpoint, which is important for OA pathophysiology and cartilage degradation, we observed that glutamine deprivation was able to decrease chondrocyte expression of inflammatory and catabolic genes in response to IL-1β stimulation. Mechanistically, we observed decreased NF-κB activation and expression of IκB-ζ with glutamine deprivation. We also note that glutamine deprivation reduces ROS generation by IL-1β stimulation, which we have previously shown is protective and can block IκB-ζ expression(Arra et al., 2022; Arra et al., 2020). This effect may be mediated via the metabolic reprogramming induced by glutamine deprivation, reversing metabolic changes induced by IL-1β that we have previously shown are pro-inflammatory. For example, LDHA and PPP activity has previously been shown to be pro-inflammatory in chondrocytes(Arra et al., 2020), but these processes are decreased in the setting of glutamine deprivation. Through this manner, glutamine deprivation may further decrease the inflammatory response through a reduction in oxidative stressors and metabolic modulation.

We then noted that glutamine deprivation is able to promote autophagy and activates stress response systems such as ATF4 pathway. These systems are likely required for maintenance of amino acid levels and anabolic activity in the absence of glutamine. However, autophagy and stress systems have also been shown to be protective and may hold therapeutic potential(Aman et al., 2021). For example, intermittent fasting and rapamycin as autophagy promoting compounds have gained interest recently as modalities for driving protective-autophagy(Johnson, Rabinovitch, & Kaeberlein, 2013; Mattson & de Cabo, 2020). Our prior work has also displayed that autophagy can also regulate inflammatory responses through modulation of NF-κB and other pathways(Adapala et al., 2020), which may be involved in the anti-inflammatory effect of glutamine deprivation. Our future work will focus on understanding the role of ATF4 stress response system and autophagy in the modulation of chondrocyte inflammatory response and NF-κB activity.

Our work then focused on the GLS reaction, which is one of the rate limiting steps of glutamine metabolism(Herranz, 2017). We displayed that ammonia is an inhibitor of autophagy and promotes inflammatory responses, while glutamate is not. Prior studies on the role of ammonia in regulation of autophagy have demonstrated that ammonia is able to both induce and inhibit autophagy through various mechanisms(Soria & Brunetti-Pierri, 2019). It is possible that ammonia derived from glutamine may be pathological, especially if it is not appropriately recycled. The role of ammonia and ammonia-removal processes in chondrocytes requires further study, especially in the context of inflammation, to determine their importance in joint disease. In addition, measurement of ammonia levels in synovial fluid may provide insight into the health of OA and RA joints. Our finding that glutamate treatment does not influence inflammatory response is in agreement with a prior study showing that exogenous glutamate did not influence inflammatory response of chondrocytes, although NMDA receptor blockade was anti-inflammatory, raising interesting questions about the role of glutamate in chondrocyte physiology(Piepoli et al., 2009). Overall, our work suggests that glutaminolysis may be pro-inflammatory through the production of ammonia which can block protective autophagy, unless systems exist for ammonia incorporation and removal that can prevent this effect.

The results of this work lays the foundation for further investigation into glutamine metabolism as a possible therapeutic target. Several inhibitors exist that may hold some therapeutic potential, such as the glutaminase inhibitor CB-839, which is currently in clinical trials for anti-tumor potential and can be repurposed for the treatment of OA. Use of genetic mouse models will also provide more detailed *in vivo* pre-clinical information when combined with OA models. In addition, further work will be performed to determine if glutamine and downstream metabolite levels can be correlated to OA disease severity, allowing for the development of biomarkers. Another major knowledge gap is the understanding of non-glutamine amino acid and fatty acid utilization by chondrocytes, which can be performed through combined metabolomic, proteomic and transcriptomics based approaches. Finally, more work needs to be performed using human samples such as articular cartilage and synovial fluid to create better translational models since OA is unlikely to be a single disease entity but a variety of sub-conditions with their own unique pathophysiology(Deveza & Loeser, 2018). A complete understanding of chondrocyte metabolism may provide an expanded toolbox for the understanding of OA and give rise to a personalized approach for patient treatment.

## Acknowledgements

This work is supported by NIH/NIAMS R01-AR072623 (to YA), Biomedical grant from Shriners Hospital for Children (YA), P30 AR074992 NIH Core Center for Musculoskeletal Biology and Medicine (to YA), R01-AR076758 and R01-AI161022 (to GM).

## Author Contributions

M.A., G.S., G.M., Y.A., participated in study design. M.A., G.S., N.S.A., K.N., L.C., performed experiments, F.R., and R.B., provided material, advice and troubleshooting, M.A., and G.S., analyzed data. M.A., and Y.A., guided the general outline and experimental approach of the project and wrote the manuscript.

## Materials and methods

### Animals

Male and female mice on C57BL/6 background were used. All the animals were housed at the Washington University School of Medicine barrier facility. All experimental protocols were carried out in accordance with the ethical guidelines approved by the Washington University School of Medicine Institutional Animal Care and Use Committee (approved protocol #21-0413).

### Murine Cell Culture

For murine chondrocyte experiments, chondrocytes were isolated from sterna of newborn pups (C57BL/6J) age P1-P3 without consideration for sex. Cells were isolated by sequential digestion with pronase (2 mg/mL, PRON-RO, Roche) at 37 degrees, followed by collagenase D (3 mg/mL, COLLD-RO, Roche) two times at 37 degrees, and cultured in DMEM (Life Technologies, Carlsbad, CA USA) containing 10% FBS and 1% penicillin/streptomycin (15140122, ThermoFisher, Waltham, MA USA) and plated in tissue culture plates. For glutamine deprivation conditions, media was changed to high glucose DMEM containing glutamine or devoid of glutamine (Life Technologies, Carlsbad, CA USA). For experiments, cells are treated with recombinant mouse IL-1β (211-11B, Peprotech, Cranbury, NJ USA) at 10 ng/mL, CB-839 (S7655, Selleck, Chemicals, Houston, TX USA), rapamycin (HY-10219, MedChem Express, Monmouth Junction, NJ USA), ammonium chloride (A9434, Sigma-Aldrich, USA), or L-glutamatic acid (G1626, Sigma-Aldrich, St. Louis, MO USA).

### Human Cell Culture

Cartilage fragments from discarded tissue post-surgery were collected in Dulbecco’s Modified Eagle Medium: Nutrient Mixture F-12 (DMEM/F-12, Gibco, ThermoFisher, Waltham, MA USA) containing 10% heat-inactivated fetal bovine serum (FBS, Gibco, ThermoFisher, Waltham, MA USA), 2% penicillin and streptomycin (10,000 U/mL, Gibco, ThermoFisher, Waltham, MA USA). Tissue fragments were digested using an enzyme cocktail containing 0.025% collagenase P (Roche, 1.5 U/mg) and 0.025% pronase (Roche, 7 U/mg)] in complete DMEM/F-12 medium in a spinner flask. After incubation at 37oC for overnight, the digest was filtered through 70 µm pore-size cell strainer and centrifuged at 1500 rpm for 5 min. Pellet was washed with calcium- and magnesium-free Hank’s Balanced Salt Solution (HBSS, Gibco, ThermoFisher, Waltham, MA USA) and suspended in complete DMEM/F-12 supplemented with 50 mg/L L-ascorbic acid.

### MLI model

Meniscal-ligamentous injury (MLI) surgery was utilized to induce OA in mice. In this procedure, medial meniscus was surgically removed from the joint without disrupting patella or other ligaments. Sham surgery was performed on the contralateral joint in which a similar incision is made on the medial side without removal of the meniscus. After 2 weeks (acute phase), mice are sacrificed and joints were collected for histology.

### Immunohistochemistry

Mouse long bones were harvested keeping knee joints intact and fixing in 10% neutral buffered formalin for 24 hours at room temperature followed by decalcification in Immunocal (StatLab, McKinney, TX) for 3 days with fresh Immunocal changed every 24 hours. Tissues were processed, embedded into paraffin, and sectioned 5 μm thick then stained with Hematoxylin-Eosin or Safranin-O to visualize cartilage and bone. For immunohistochemistry, sections were deparaffinized and rehydrated using 3 changes of xylenes followed by ethanol gradient. Antigen retrieval in murine sections was performed by incubating samples in Citrate buffer (pH 6.0) at 55 degrees Celsius overnight, followed by washing in PBS and subsequent quenching of endogenous peroxidase activity by incubation in 3% H2O2 for 15 minutes at room temperature. Sections were then blocked using blocking solution (10% normal goat serum, 5% BSA, 0.1% Tween-20) for 1 hour at room temperature. Sections were incubated overnight at 4 degrees with anti-ATF4 (10835-1-AP, ThermoFisher, Waltham, MA USA, RRID: AB_2058600) antibody at a 1:200 dilution. Sections were rinsed in phosphate-buffered saline (PBS) several times followed by addition of 1:500 dilution of biotinylated secondary antibody (BP-1100, Vector Biolabs, Burlingham, CA) for one hour. Post-secondary antibody incubation, sections were washed with PBS-T several times followed by incubation with streptavidin-HRP (2 μg/ml) for 20 min. After extensive washing with PBS, sections were developed using DAB peroxidase kit (SK4100, Vector Biolabs, Burlingham, CA), with development on each slide standardized to the same amount of time.

### Protein analysis by Immunoblotting

Cell lysates for protein analysis were prepared by scraping cells in 1x Cell Lysis Buffer (Cell Signaling Technology, Danvers, MA, USA) containing 1x protease/phosphatase inhibitor (Thermo Fisher Scientific, Waltham, MA USA Halt Protease Phosphatase Inhibitor Cocktail). Blotting was performed using primary antibodies for LC3B (2775, CST, Danvers, MA USA, RRID:AB_915950), p62 (2C11, Abnova, Taiwan, RRID:AB_437085), IκB-ζ (Cat# 14-16801-82, Invitrogen, RRID:AB_11218083), p-AKT (9271, CST, Danvers, MA USA RRID: AB_329825), total Akt (9272, CST, Danvers, MA USA, AB_329827), phospho-S6 (2211, CST, Danvers, MA USA, RRID:AB_331679), total S6 (2217, CST, Danvers, MA USA, RRID:AB_331355) and Actin (Cat# A2228, Sigma, St. Louis, MO RRID:AB_476697). Protein concentration was determined by BCA assay (23225, Pierce, ThermoFisher, Waltham, MA USA) and equal amounts of protein were loaded onto SDS-PAGE gel.

### Immunocytochemistry

Chondrocytes were plated on sterile, gelatin-coated glass coverslips placed in 24 well plates at lower concentration. Cells were cultured under normal media conditions and treatments were performed in the 24 well plate. For staining, media was removed and cells were fixed in 4% formaldehyde in PBS for 30 minutes. Cells were washed with PBS containing 0.1% saponin. Cells were blocked using blocking buffer (1x PBS, 5% NGS, 0.1% saponin) for 1 hour at room temperature. Cells were incubated with anti-LC3b (12741, CST, Danvers, MA USA RRID:AB_2617131) or anti-p62 (2C11, Abnova, Taiwan, RRID:AB_437085) antibodies at 1:100 concentration in antibody dilution buffer (1x PBS, 1% BSA, 0.1% saponin) overnight at 4 degrees. Cells were washed with wash buffer (1x PBS, 0.1% saponin) 3 times and incubated with fluorescent conjugated secondary antibody at 1:1000 in antibody dilution buffer for 2 hours at room temperature. Samples were washed with wash buffer 3 times. Slides were coverslipped with antifade mounting media containing DAPI (9071, CST, Danvers, MA USA). Images were taken on fluorescent microscope.

### Measurement of extracellular lactic acid

Chondrocytes were cultured for 1 day with IL-1β treatment (10 ng/mL) with appropriate experimental conditions in 96 well plates containing 200 μl of DMEM containing 10% FBS. Supernatant media was collected and centrifuged to separate cell debris and floating cells. Supernatant was used immediately for lactic acid assay to measure secreted lactate in the media using a 1:20 dilution (Cat# 1200011002, Eton Biosciences, San Diego, CA USA). Unconditioned DMEM with 10% FBS was used as a control for subtracting background.

### Measurement of gene expression by qPCR

Trizol (Sigma, St. Louis, MO USA) was added to samples to isolate mRNA from cell culture samples. Chloroform was added at a ratio of 0.2:1 to Trizol to samples, followed by centrifugation at 12,000g for 15 mins. Aqueous layer was isolation and equal amount of 70% ethanol was added. RNA was then isolated from this fraction using PureLink RNA mini kit (Cat# 12183025, Ambion, Grand Island, NY, USA) and cDNA was prepared using High Capacity cDNA Reverse Transcription kit (Cat# 4368814, Applied Biosystems). qPCR was carried out on BioRad CFX96 real time system using iTaq universal SYBR green super-mix (Cat#1725120, BioRad, Hercules, CA, USA). mRNA expression was normalized using actin as a housekeeping gene. Full list of primers is listed in Table 1.

**Table 1:**
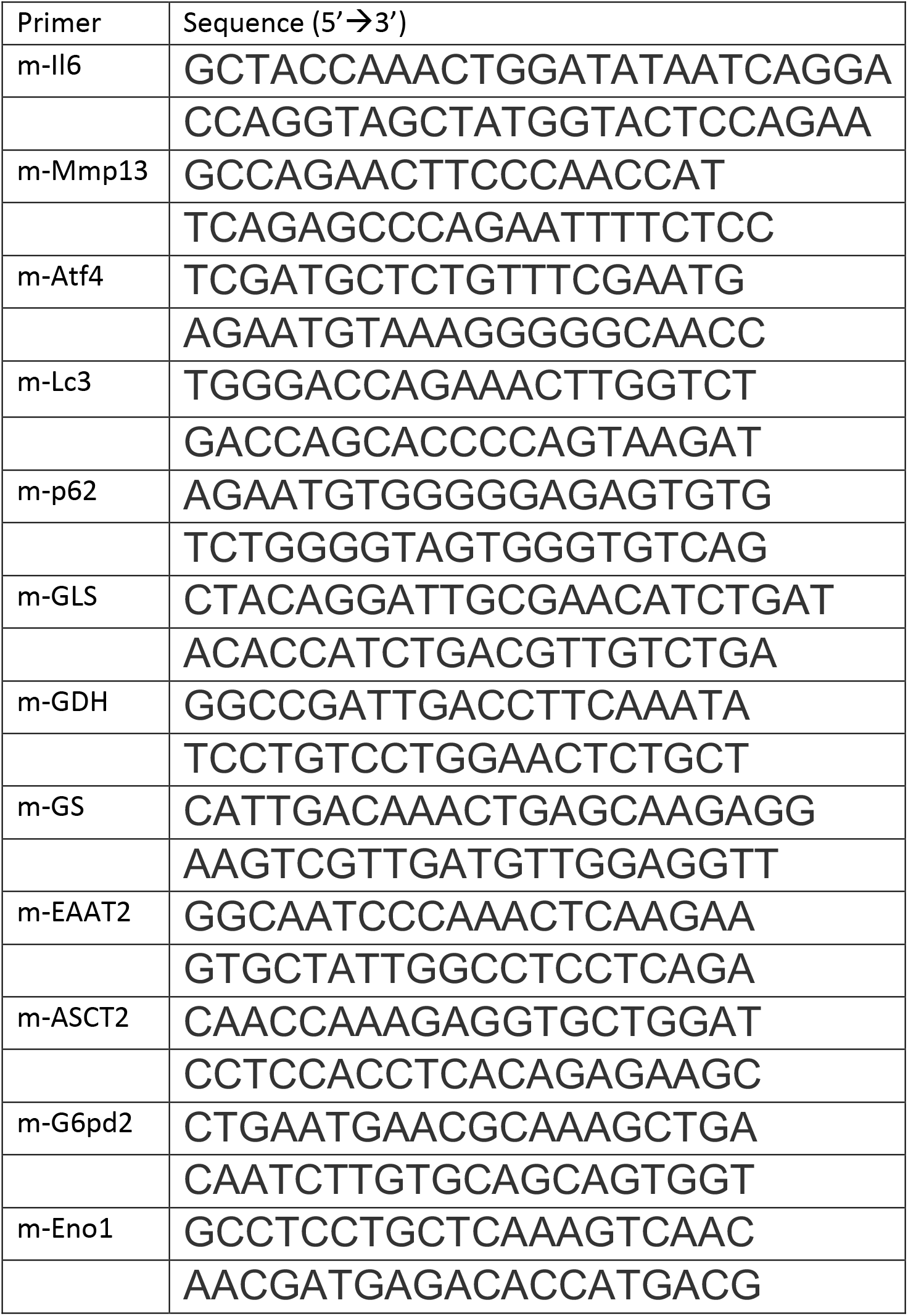
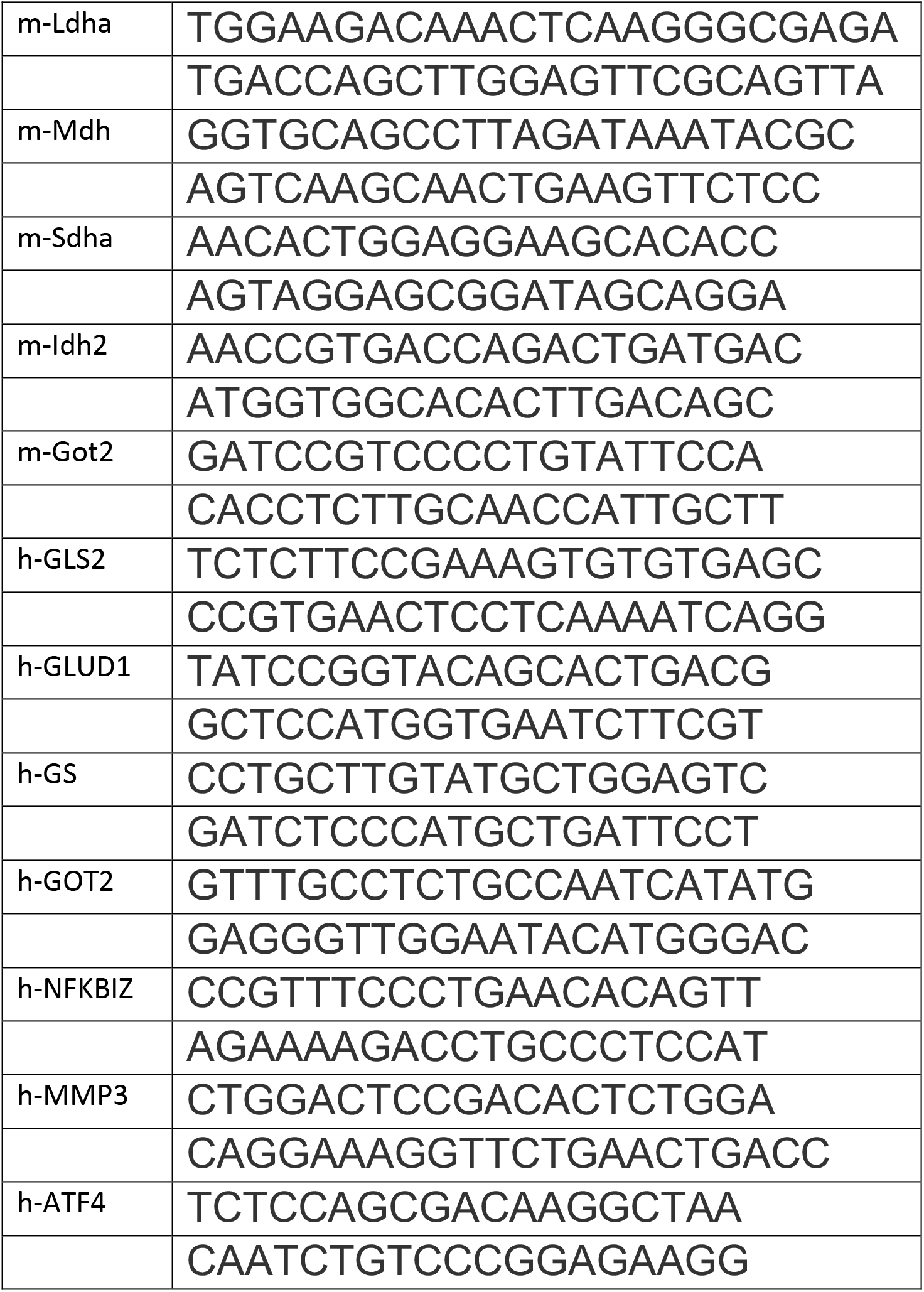
List of primers:

### Measurement of cellular metabolism by Seahorse

Primary chondrocytes were plated in Seahorse XF96 plates at 50,000 cells per well and cultured in media containing glutamine or without glutamine. Cells were then treated with IL-1β (10 ng/mL). After 24 hours, Seahorse assay was performed. For glycolysis stress test, cells were serum starved for 1 hour in glucose-free media containing treatments, and measurement of ECAR and OCR was performed prior to and after sequential addition of glucose, oligomycin and 2-DG with measurements performed every 5 minutes. For MitoStress test, cells were incubated in glucose-containing media for 1 hour containing treatments and measurements were performed every 5 minutes prior to and after sequential addition of oligomycin, FCCP and Rotenone/Antimycin A. Media for seahorse assays was devoid of glutamine. Data was analyzed using Wave software.

### Measurement of intracellular ATP

Primary chondrocytes were plated in 96 well plates at 5x10^4^ and treated with IL-1β for 24 hours. Lysates were collected and processed according to luminescence-based ATP assay kit (Cat#K255, Biovision, Milpitas, CA USA ADP/ATP ratio Assay kit). Assay was performed in 96 well plate. Luminescence was measured using luminescent plate reader after 15 minutes. Data was collected and processed using Gen5 software.

### Measurement of ROS

Primary chondrocytes were treated for 24 hours in DMEM media. Cells were washed two times with phenol red-free PBS, followed by incubation with 10 μM DCFDA (Cat#D6883, Sigma, St. Louis, MO USA) in PBS for 30 minutes, followed by two more washes with PBS. Cells were incubated in 37 degrees C incubator for one hour in PBS, followed by fluorescence measurement using microplate reader using Ex/Em 495/525 for DCFDA.

### Measurement of metabolite concentrations

The cell suspensions (2 x 10^6^ cells/mL) were prepared by vortexing cell pellets with water. The amino acids and metabolites listed above were extracted from 50 µL of cell suspension with 200 µL of methanol after addition of internal standards (Glu-d3 (1.6 µg), Asp-d3 (1.6 µg), Asn-d3,15N2 (1.6 µg), Gln-13C5 (1.6 µg), alpha-ketoglutarate-d2 (0.4 µg), 2-hydroxyglutarate-d3 (0.2 µg), oxaloacetate-13C3 (0.2 µg), pyruvate-13C3 (2 µg), and malate-d3 (0.2 µg)). The sample aliquots for alpha-ketoglutarate, oxaloacetate, and pyruvate were derivatized with o-phenylenediamine to improve mass spectrometric sensitivity. Quality control (QC) samples were prepared by pooling aliquots of study samples to monitor instrument performances throughout these analyses.

The analysis of Gln, Glu, Asp, and Asn was performed on a Shimadzu 20AD HPLC system and a SIL-20AC autosampler coupled to 4000Qtrap mass spectrometer (Applied Biosystems) operated in positive multiple reaction monitoring (MRM) mode. The analysis of 2-hydroxyglutarate and malate was performed on a Shimadzu 20AD HPLC system and a SIL-20AC autosampler coupled to 4000Qtrap mass spectrometer (Applied Biosystems) operated in negative MRM mode. The analysis of alpha-ketoglutarate, oxaloacetate, and pyruvate was performed in positive ion MRM mode on API4000 mass spectrometer (Applied Biosystems) coupled to a Shimadzu 20AD HPLC system and a SIL-20AC autosampler. The QC samples were injected every five study samples. Data processing was conducted with Analyst 1.6.3 (Applied Biosystems).

### Measurement of viability

Cells were treated under appropriate conditions. MTT ((3-(4,5-dimethylthiazol-2-yl)-2,5-diphenyltetrazolium bromide) was dissolved in PBS at 10 mg/mL. MTT was added to wells to a final concentration of 0.5 mg/mL. Cells were incubated at 37 degrees for 6 hours. Media was removed and 50 uL of DMSO was added to each well. Plate was placed on a plate shaker and measurement was made at 580 nm on microplate reader. Data was collected and processed using Gen5 software.

### Measurement of intracellular glutamate

Intracellular glutamate was measured in chondrocytes in 96 well plate format utilizing Glutamate-Glo assay kit (J7021, Promega, Madison, WI USA). Luminescence was measured on microplate reader. Data was collected and processed using Gen5 software.

### NF-κB Luciferase Assay

Chondrocytes were isolated from sterna of newborn NF-κB-luciferase reporter mice as described earlier. Cells were cultured and treated appropriately in 96 well plate tissue culture plate. Cells were washed twice with 1x PBS and lysed using 1x luciferase lysis buffer (L-740, GoldBio, St. Louis, MO USA). Plates were freeze-thawed in -80 freezer. 20 uL of lysates were transferred to white bottom, round well microplates. Detection was performed after addition of 50 uL of detection reagent (I-930, GoldBio, St. Louis, MO USA). Luminescence was measured by microplate reader. Data was collected and processed using Gen5 software.

### Statistical Analysis

All experiments represent biological replicates and were repeated at least three times, unless otherwise stated. Technical replicates are considered to be repeat tests of the same value, i.e. testing same samples in triplicate for qPCR. Biological replicates are considered to be samples derived from separate sources, such as different mice or on different dates. Statistical analyses were performed using appropriate statistical test using GraphPad Prism. All graphs were generated using Prism as well. Multiple treatments were analyzed by One-way ANOVA followed by Tukey’s test multiple comparisons test for greater than 2 groups. Student’s T test was used for comparing two groups. Student’s T test was performed for comparing same biological samples subject to different treatments. *P* values are indicated where applicable. *P<0.05, **P<0.01, ***P<0.0005, ****P <.0001. Histology and immunostaining data were scored by investigators blinded to the experimental conditions. Male and female mice were used at equal ratios for cell culture to avoid sex bias. Sample size determination was not required.

## Figures

**Figure 1 Supplement 1.**
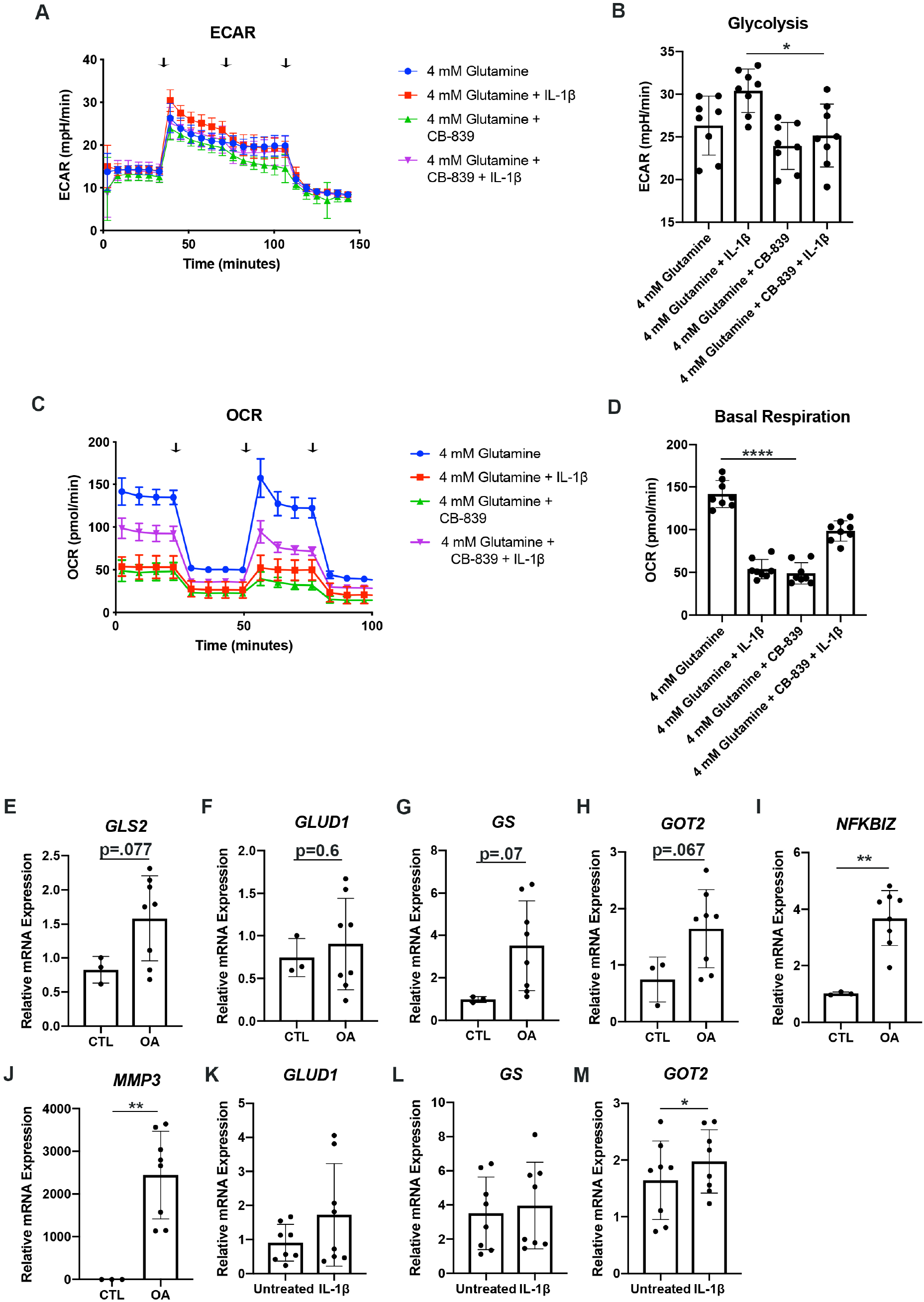
**(A-D)** Primary sternal chondrocytes were cultured in media containing glutamine with or without CB-839 (1 mM). Cells were then treated with IL-1β (10 ng/mL) for 24 hours. All values were normalized to cell viability of treatments relative to untreated cells as measured by MTT assay. (**A-B**) ECAR measurement in glycolysis stress test (Injection 1: No treatment, Injection 2: Glucose, Injection 3: Oligomycin, Injection 4: 2-DG) or (**C-D**) OCR measurement in MitoStress test (Injection 1: No treatment, Injection 2: Oligomycin, Injection 3: FCCP, Injection 4: Antimycin A/Rotenone) was performed on Seahorse Instrument. Measurements were performed every 6 minutes with 8 replicates per timepoint for each condition. Arrows represent injection timepoints. Graphs shown in Fig S1B and S1D are from a single timepoint. One-way ANOVA was performed followed by Tukey’s multiple comparisons test. B: *P= 0.012, D: ****P<0.0001. **(E-J)** Human chondrocytes were isolated from OA cartilage or healthy cartilage isolated from patients. Gene expression was measured by qPCR. **(K-M)** Human chondrocytes were isolated from knee cartilage and cultured in media. Cells were treated with IL-1β for 24 hours. Gene expression was measured by qPCR from n=8 biological samples. Unpaired student’s T test was performed. I:**P=0.0013, J:**P=0.0032, M:*P=0.05.

**Figure 2 Supplement 2.**
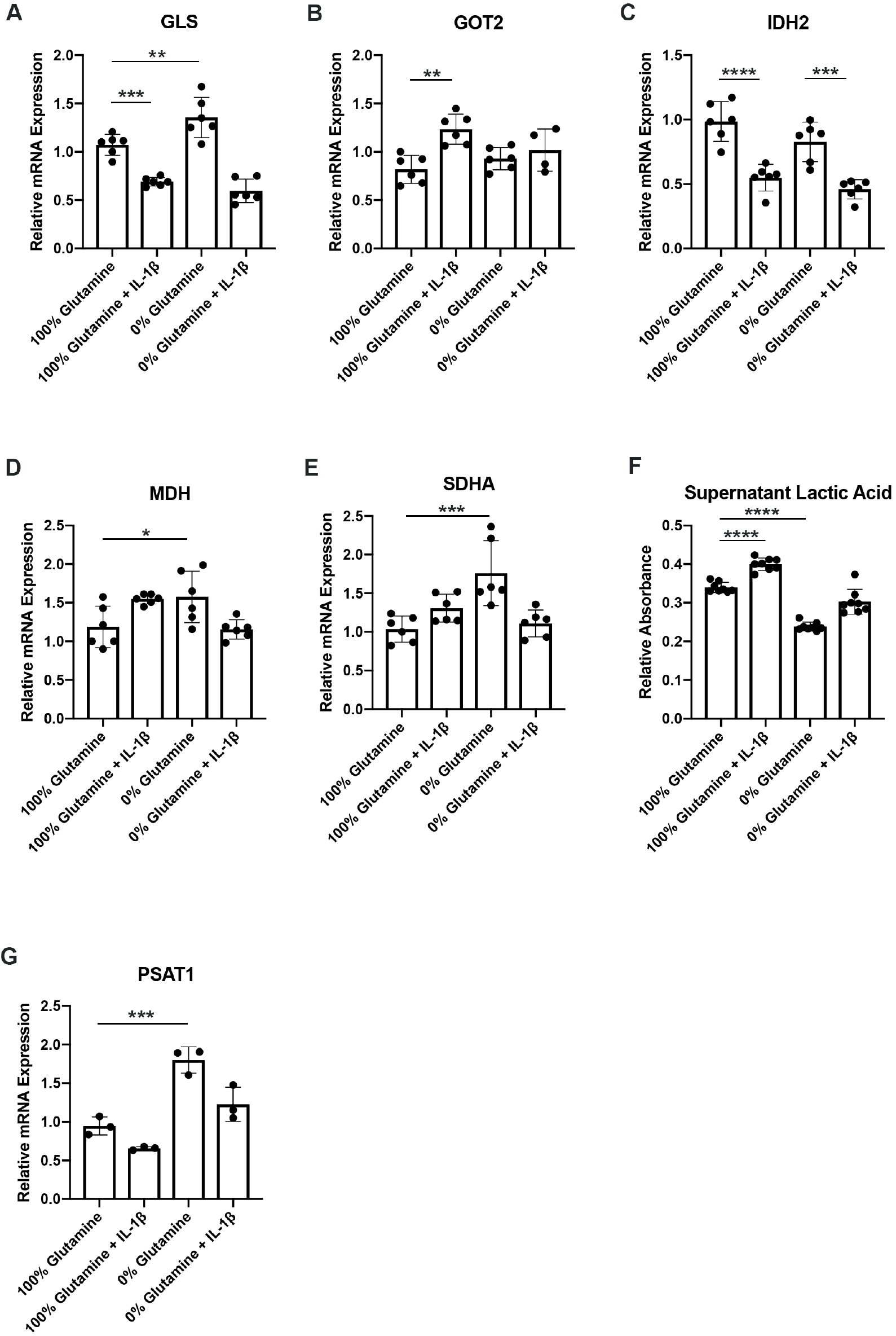
**(A-E)** Primary murine chondrocytes were cultured in media containing 100% glutamine or 0% glutamine for 24 hours. Cells were then treated with IL-1β (10 ng/mL) for 24 hours. Gene expression of *Gls*, *Got2*, *Idh2, Mdh, and Sdha* were measured by qPCR from n=6 replicates. One-way ANOVA was performed followed by Tukey’s multiple comparisons test. A: **P=0.0085, ***P<0.0004, B:**P=0.0011, C: ****P<0.0001, ***P=0.0003, D:*P=0.036, E:***P=0.0005. **(F)** Under similar conditions, supernatant was collected and lactic acid was measured. One-way ANOVA was performed followed by Tukey’s multiple comparisons test. ****P<0.0001. **(G)** Gene expression of *Psat1* was measured under similar conditions by qPCR. One-way ANOVA was performed followed by Tukey’s multiple comparisons test. ***P=0.0006.

**Figure 3 Supplement 3.**
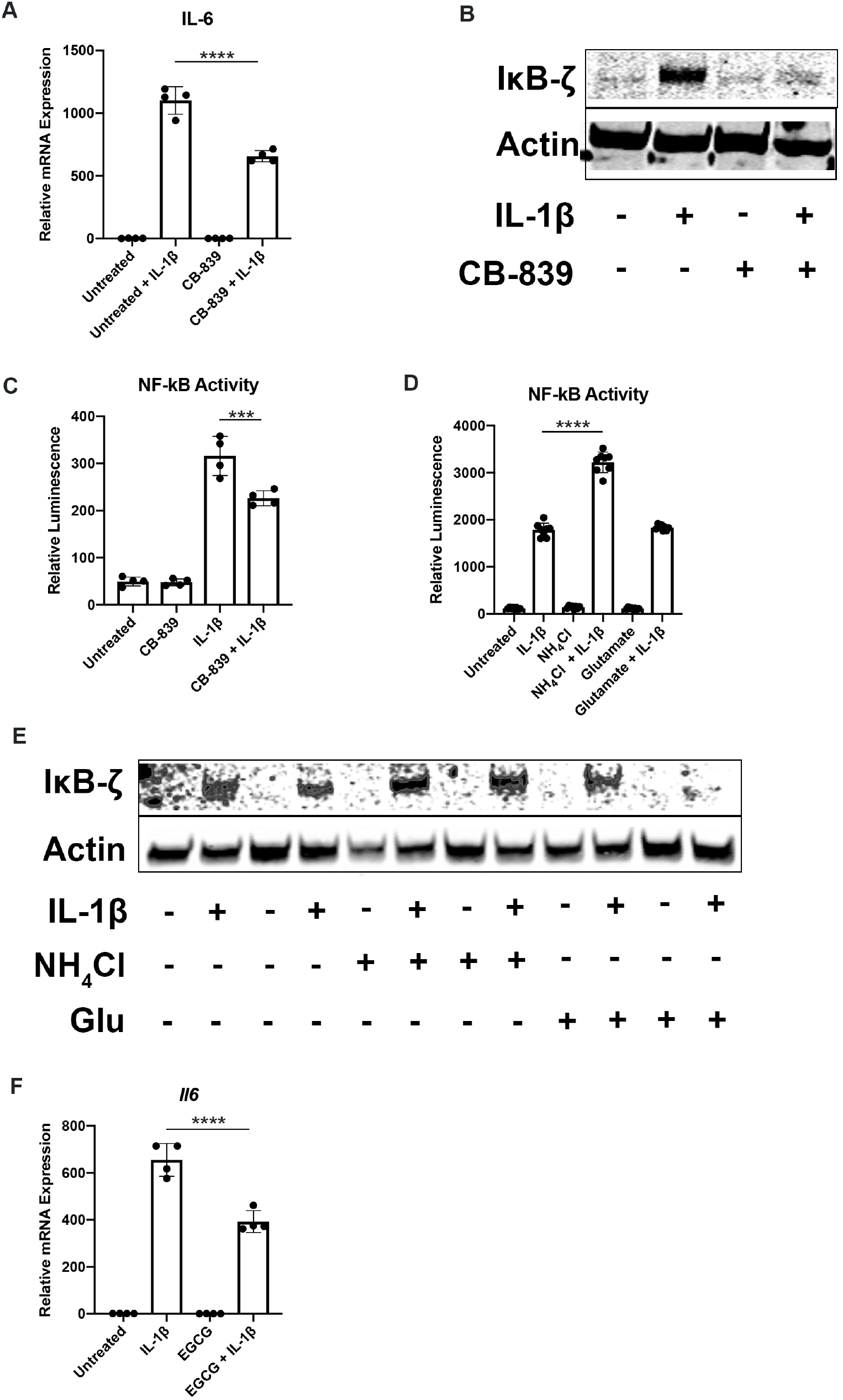
**(A)** Primary murine chondrocytes were treated with IL-1β in the presence or absence of CB-839 (1 mM) for 24 hours. Gene expression of *Il6* was measured by qPCR. Results are representative of one experiment. One-way ANOVA was performed followed by Tukey’s multiple comparisons test. ****P<0.0001. **(B)** Under similar conditions, IκB-ζ protein levels were measured by immunoblotting. **(C)** Primary NF-κB-luciferase reporter chondrocytes were treated with IL-1β in the presence or absence of CB-839 (1 mM) for 24 hours. NF-κB activity was measured by luciferase assay. One-way ANOVA was performed followed by Tukey’s multiple comparisons test. ***P=0.0007 **(D)** Primary NF-κB-luciferase reporter chondrocytes were treated with IL-1β in the presence or absence of ammonium chloride (2 mM) or glutamate (200 μM). NF-κB activity was measured by luminescent luciferase assay. One-way ANOVA was performed followed by Tukey’s multiple comparisons test. ****P<0.0001. **(E)** Under similar conditions, lysates were collected and IκB-ζ protein levels were measured by immunoblotting. Image from representative experiment. **(F)** Chondrocytes were treated with IL-1β in the presence or absence of ECGC for 24 hours. Gene expression of *Il6* was measured by qPCR. Results are representative of one experiment. One-way ANOVA was performed followed by Tukey’s multiple comparisons test. ****P<0.0001.

**Figure 4 Supplement 4.**
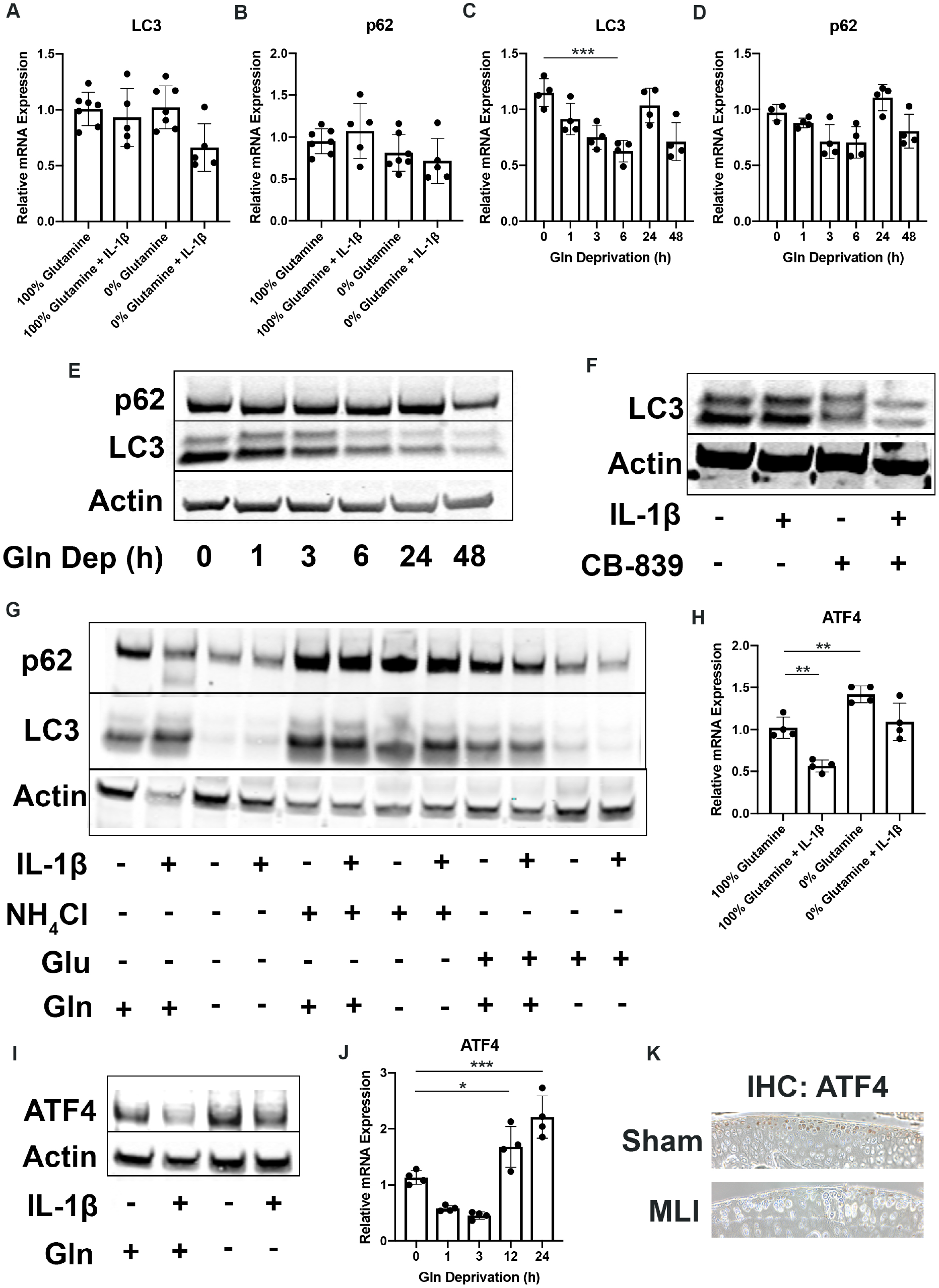
**(A-B)** Primary murine chondrocytes were cultured in media containing 100% glutamine or 0% glutamine for 24 hours. Cells were then treated with IL-1β (10 ng/mL) for 24 hours. Gene expression of *Lc3* and *p62* was measured by qPCR. Results from 5-6 biological replicates. One-way ANOVA was performed followed by Tukey’s multiple comparisons test. **(C-D)** Chondrocytes were plated in glutamine-free media for different amounts of time. Gene expression of *Lc3* and *p62* was measured by qPCR. One-way ANOVA was performed followed by Tukey’s multiple comparisons test. ***P=0.0004. **(E)** Under similar conditions, immunoblotting was performed for LC3 and p62. **(F)** Chondrocytes were treated with IL-1β in the presence or absence of CB-839 for 24 hours. Immunoblotting was performed for LC3. **(G)** Primary chondrocytes were cultured in media containing 4 mM or 0 mM glutamine. Cells were supplemented with ammonium chloride (2 mM) or glutamate (200 μM). After 6 hours, cells were treated with IL-1β (10 ng/mL) for 24 hours. Immunoblotting was performed for p62 and LC3B. Image displays representative experiment. **(H)** Primary murine chondrocytes were cultured in media containing 100% glutamine or 0% glutamine for 24 hours. Cells were then treated with IL-1β (10 ng/mL) for 24 hours. Gene expression of *Atf4* was measured by qPCR. Results from n=4 replicates. One-way ANOVA was performed followed by Tukey’s multiple comparisons test. **P=0.0033, **P=0.0088. **(I)** Immunoblotting was performed under similar conditions for ATF4. **(J)** Chondrocytes were plated in glutamine-free media for different amounts of time. Gene expression of *Atf4* was measured by qPCR. One-way ANOVA was performed followed by Tukey’s multiple comparisons test. *P=0.0419, ***P=0.0001. **(K)** MLI surgery was performed on 12 week old mice to induce OA with sham surgery performed on contralateral leg. After 2 weeks, knee joints were collected and sectioned. IHC was performed for ATF4 which is display as brown stain, with representative image displayed.

**Figure 5 Supplement 5.**
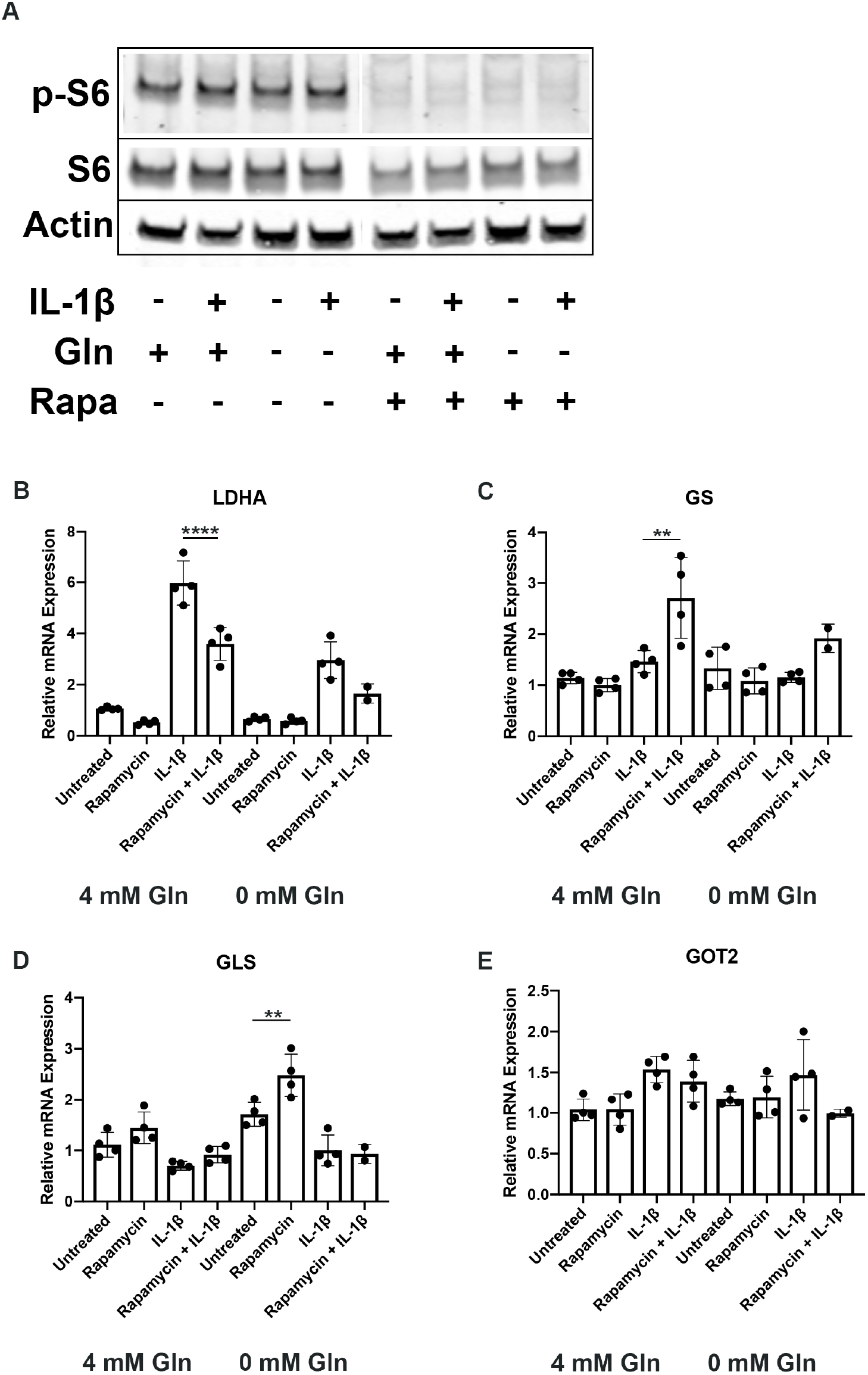
**(A)** Primary murine chondrocytes were cultured in media containing 100% glutamine or 0% glutamine for 24 hours. Cells were then treated with IL-1β (10 ng/mL) for 24 hours in the presence or absence of rapamycin (50 nM). Immunoblotting was performed for pS6 and total S6. **(B-E)** Under similar conditions, gene expression was measured by qPCR. One-way ANOVA was performed followed by Tukey’s multiple comparisons test. B:****P<0.0001, C: **P=0.0017, D:**P=0.0088.

